# A Dual-Compartment Scaffolding Role for RACK1 in Hepatic Glucagon Signaling and Gluconeogenesis

**DOI:** 10.1101/2025.06.18.660434

**Authors:** Cancan Lyu, Ling Yang, Songhai Chen

## Abstract

**Background & Aims:** The hepatic glucagon–PKA–CREB signaling axis plays a central role in regulating gluconeogenesis and maintaining glucose homeostasis during fasting. However, the mechanisms that govern the spatial coordination and substrate specificity of this pathway remain incompletely understood. This study determines the role of the scaffolding protein RACK1 (Receptor for Activated C Kinase 1) in orchestrating glucagon signaling to regulate hepatic gluconeogenesis.

**Methods:** RACK1 was acutely deleted in mouse liver and primary hepatocytes. Metabolic phenotypes were assessed by glucose, pyruvate, and insulin tolerance tests, and hepatocyte glucose production assays. Protein interactions were analyzed with co-immunoprecipitation, GST pulldown, and proximity ligation assays. Subcellular localization and signaling events were studied by Western blot, confocal microscopy, and fractionation. Functional rescue was performed by hepatic expression of a constitutively active PKA catalytic subunit (PKAcα^W196R^).

**Results:** Acute hepatic RACK1 deficiency caused fasting hypoglycemia, impaired gluconeogenesis, and improved glucose and pyruvate tolerance without affecting insulin signaling. RACK1 directly bound GCGR, PKA regulatory (RIIα) and catalytic (PKAcα) subunits, and CREB, functioning as a dual-compartment scaffold assembling GCGR–PKA complexes at the plasma membrane and PKAcα–CREB complexes in the nucleus. Loss of RACK1 impaired PKAcα translocation, CREB phosphorylation, and gluconeogenic gene expression. These defects were rescued by PKAcα^W196R^ expresson. Overexpression of RACK1 WD1–2 and WD3–4 domains, which mediate PKA and GCGR interactions, similarly disrupted PKA signaling and gluconeogenesis.

**Conclusion:** RACK1 spatially organizes the glucagon–PKA–CREB axis, ensuring precise signal propagation and efficient hepatic gluconeogenesis, revealing a novel mechanism of compartmentalized signal regulation in glucose metabolism.

## INTRODUCTION

Maintaining glucose homeostasis is critical for energy balance and optimal cellular function [1]. The liver plays a central role in regulating systemic glucose levels [2]. Following glucose intake, the liver stores excess glucose as glycogen. During fasting, it releases glucose into the bloodstream to maintain normoglycemia and meet energy demands, particularly for the brain, through glycogenolysis and gluconeogenesis.

Glucagon signaling plays a central role in hepatic glucose metabolism, regulating both gluconeogenesis and glycogenolysis [1, 3]. In response to hypoglycemia, glucagon secreted by the pancreas binds to its receptor on hepatocytes, activating G-protein signaling predominantly via Gs proteins [4]. This activation triggers adenylyl cyclase to produce cyclic AMP (cAMP), which subsequently activates protein kinase A (PKA). PKA phosphorylates key metabolic enzymes, such as phosphorylase kinase, pyruvate kinase, and phosphofructokinase 2, to acutely regulate glucose production [5]. Furthermore, PKA phosphorylates transcription factors like CREB (cAMP response element-binding protein), which drives the expression of gluconeogenic genes, including glucose-6-phosphatase catalytic subunit (G6PC) and phosphoenolpyruvate carboxykinase (PEPCK) [2]. These mechanisms ensure adequate glucose production to meet the body’s energy demands during fasting. Dysregulation of the glucagon-PKA signaling pathway can result in metabolic disorders, such as hyperglycemia in diabetes [6, 7], highlighting the critical need for precise regulation of this pathway to maintain metabolic health.

The PKA holoenzyme consists of two regulatory subunits and two catalytic subunits (PKAc) [8]. PKA activation is initiated by cAMP binding to regulatory subunits, releasing active PKAc subunits. PKA activity is tightly regulated spatially and temporally by A-kinase anchoring proteins (AKAPs), which tether PKA to specific subcellular locations and assemble other signaling proteins to ensure specificity [9]. Over 50 AKAPs have been identified, with at least 10 expressed in the liver [9, 10]. However, recent studies suggest that most liver-expressed AKAPs, except radixin, do not significantly influence glucagon-induced PKA activation and gluconeogenesis [10]. It remains unknow if there are other non-AKAP proteins involved in regulating PKA signaling and gluconeogenesis. This gap highlights the need for further research into how the glucagon-PKA signaling axis is regulated to control glucose metabolism effectively.

RACK1 is a scaffolding protein that coordinates multiple signaling pathways and cellular functions [10]. It has been implicated in ischemia-reperfusion-induced liver injury and the development of liver cancer [11]. Recent studies also highlight a role for RACK1 in hepatic glucose and lipid metabolism. Liver-specific deletion of RACK1 in fetal mice using Alb-Cre (Cre recombinase driven by the albumin promoter and enhancer) [12] leads to progressive hepatic steatosis and tumorigenesis, likely due to impaired autophagy initiation [13]. These mice also exhibit fasting-induced hypoglycemia and improved glucose tolerance, suggesting a role for RACK1 in regulating hepatic glucose homeostasis [14]. However, because RACK1 was deleted during embryogenesis, it remains unclear whether these metabolic effects reflect a direct role of RACK1 or result from compensatory adaptations to other defects. Furthermore, the molecular mechanisms through which RACK1 modulates glucose metabolism have yet to be elucidated.

We investigated RACK1’s role in glucose metabolism by acutely deleting it in mouse liver and hepatocytes. RACK1 deficiency caused fasting hypoglycemia, impaired glucose homeostasis, and reduced gluconeogenesis. Unlike AKAPs, RACK1 binds both PKA regulatory and catalytic subunits, as well as GCGR and CREB, facilitating PKA’s membrane localization near GCGR and nuclear translocation for CREB activation. Expression of a constitutively active PKA catalytic mutant rescued these defects.

These findings reveal RACK1 as a previously unrecognized critical scaffold coordinating glucagon–PKA–CREB signaling across compartments to regulate hepatic gluconeogenesis and maintain glucose homeostasis.

## RESULTS

### Acute RACK1 deficiency results in hypoglycemia, impaired glucose homeostasis, and reduced hepatic gluconeogenesis

To investigate the direct effect of RACK1 deficiency on glucose metabolism, we acutely deleted the RACK1 gene in the liver by injecting AAV vectors encoding Cre recombinase under the control of the TBG promoter (AAV-TBG-iCre) into the retro-orbital vein of RACK1^fl/fl^ transgenic mice. Glucose metabolism was assessed 10 days post-injection. Acute hepatic RACK1 deficiency significantly reduced blood glucose levels after 6 to 18 hours of fasting, while blood glucose levels in the fed state remained unaffected (Figure 1A). Insulin, glucose, and pyruvate tolerance tests revealed that acute hepatic RACK1 deficiency significantly improved glucose and pyruvate tolerance without affecting insulin tolerance (Figure 1B-1F). These findings indicate that hepatic RACK1 deficiency enhances glucose and pyruvate tolerance by suppressing gluconeogenesis, without altering systemic insulin sensitivity.

**Figure 1.**
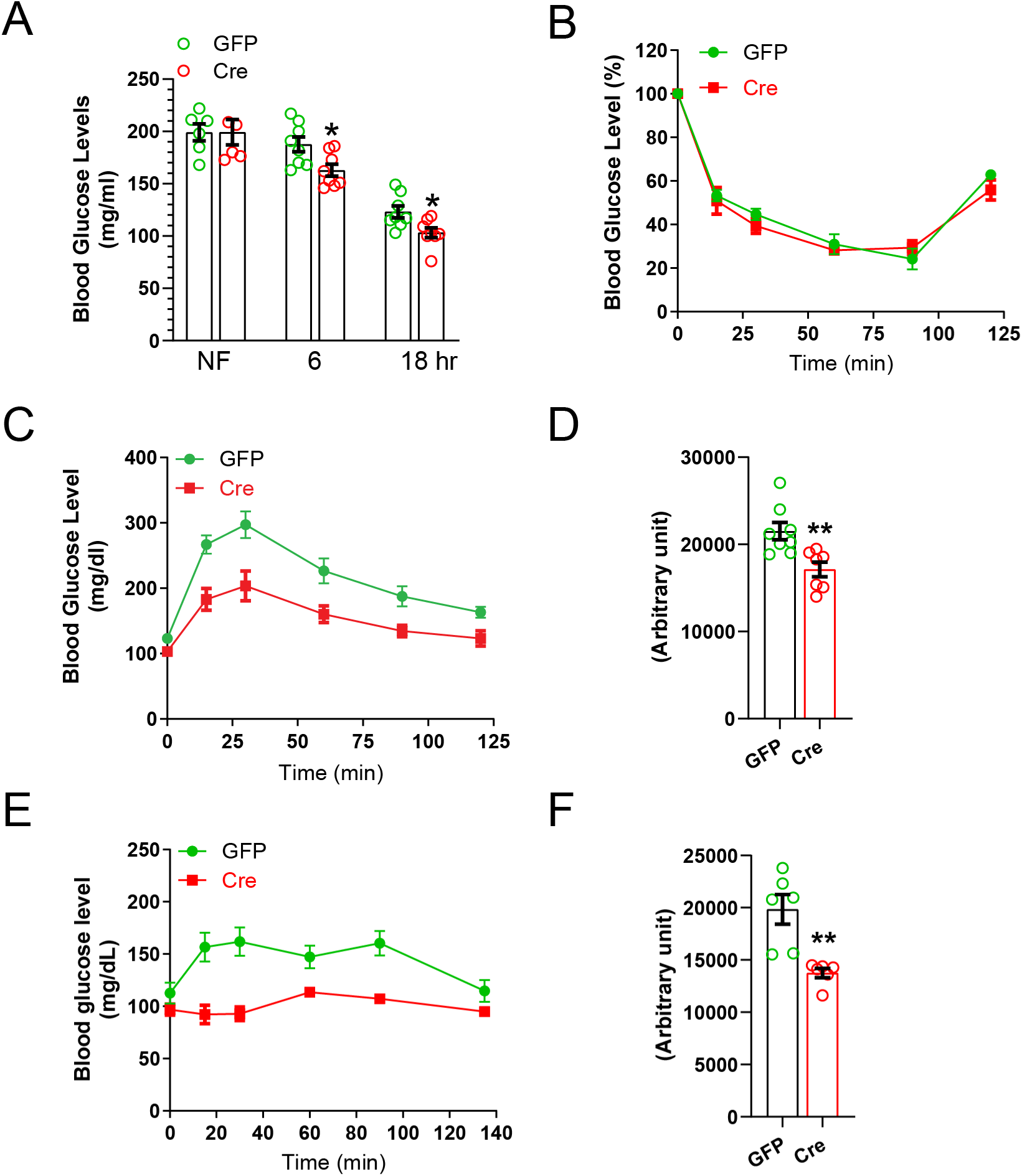
Acute RACK1 deficiency disrupts glucose homeostasis. **(A)** Blood glucose levels in RACK1^fl/fl^ mice injected with AAV8-TBG-GFP (GFP) or AAV8- TBG-iCre (Cre), measured under non-fasted (NF) conditions and after 6 or 18 hours of fasting. **(B–F)** Metabolic tolerance tests in GFP and Cre mice: **(B)** insulin tolerance test (ITT), **(C–D)** glucose tolerance test (GTT), and **(E–F)** pyruvate tolerance test (PTT). **(D)** and **(F)** show area under the curve (AUC) quantification for (C) and (E), respectively. *, **, ***p < 0.05, 0.01 and 0.001 compared to GFP controls, respectively.

Supporting this notion, glucose production assays in primary hepatocytes demonstrated that RACK1 deficiency impaired glucagon-induced gluconeogenesis but did not affect insulin-mediated suppression of glucose production (Figure 2A).

**Figure 2:**
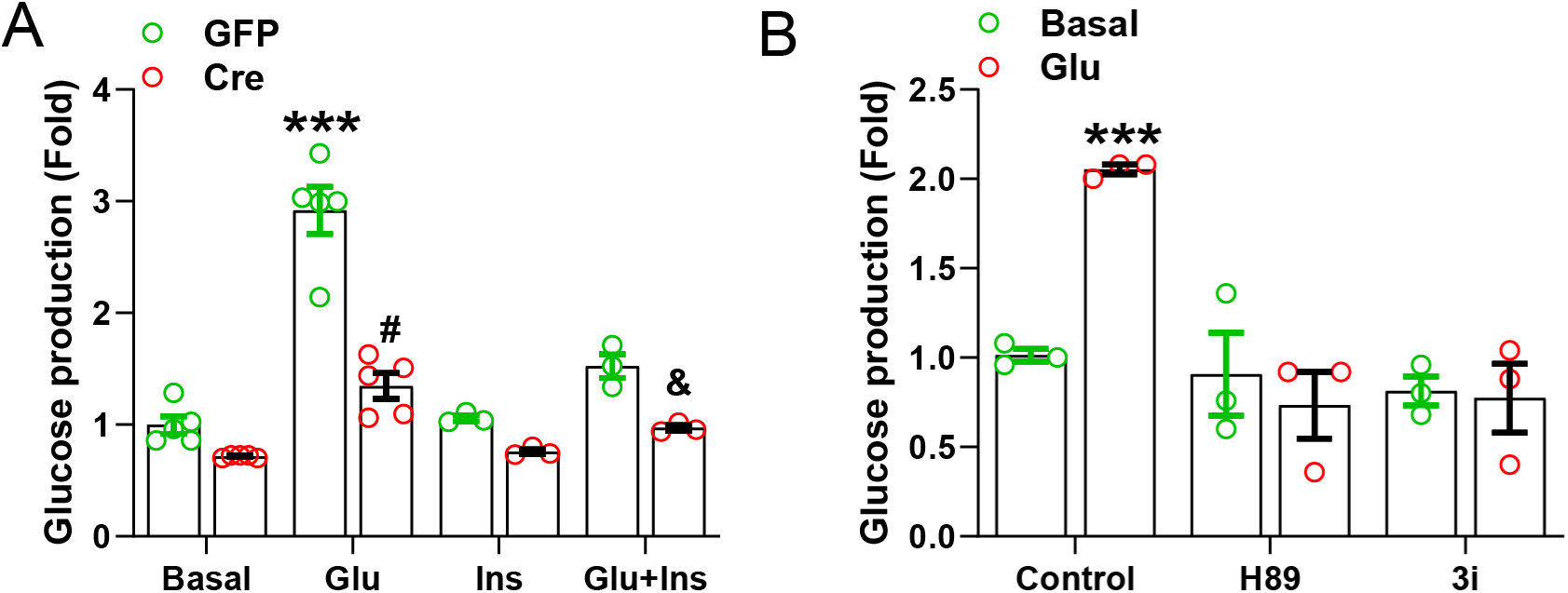
**RACK1 deficiency impairs hepatic gluconeogenesis *in vitro.*** Primary hepatocytes were isolated from RACK1^fl/fl^ mice injected with AAV8-TBG-GFP (GFP) or iCre (Cre) and subjected to glucose production assays. **(A)** Glucose output was measured under basal conditions or following stimulation with glucagon (Glu, 200 nM), insulin (Ins, 20 nM), or both. **(B)** Cells were treated with vehicle (control) or the PKA inhibitors H89 (5 μM) and compound 3i (0.5 μM) in the absence or presence of glucagon (200 nM). ***p < 0.001 vs. GFP basal; ^#^ and ^&^p < 0.05 vs. GFP.

### RACK1 selectively regulates PKA signaling

Glucagon stimulates gluconeogenesis through the PKA-CREB signaling axis [11]. Consistent with this mechanism, treatment of primary hepatocytes with PKA and CREB inhibitors (H89 and 3i, respectively) effectively blocked glucagon-induced glucose production (Figure 2B). These findings suggest that RACK1 may influence the glucagon-PKA-CREB pathway to regulate hepatic gluconeogenesis.

To test this hypothesis, we first assessed whether RACK1 deficiency affects cAMP production in the livers of fasted mice. As shown in Figure 3A, cAMP levels were comparable between RACK1^fl/fl^ mice injected with AAV8-TBG-GFP or AAV8-TBG-iCre, indicating that RACK1 does not regulate cAMP generation. However, phosphorylation of CREB at Ser133, pCREB^S133^, was significantly reduced in RACK1^fl/fl^ mice injected with AAV8-TBG-iCre compared to AAV8-TBG-GFP, whereas phosphorylation of AKT at Ser473 (pAKT^^473^) remained unaffected (Figure 3B-C). These results suggest that RACK1 specifically modulates glucagon signaling by acting on PKA rather than insulin signaling.

**Figure 3:**
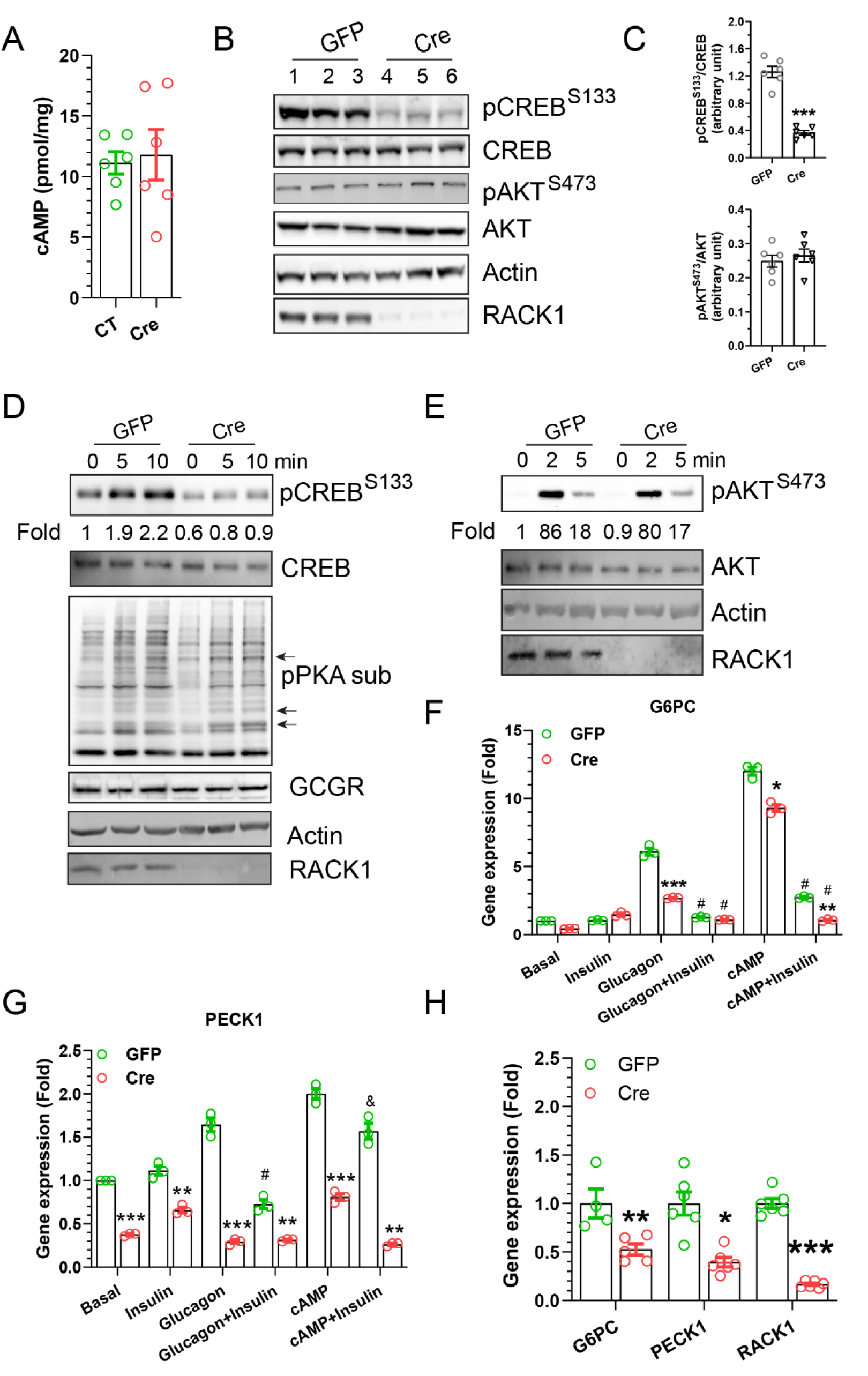
RACK1 deficiency attenuates hepatic PKA signaling**. (A)** Hepatic cAMP levels in RACK1^fl/fl^ mice injected with AAV8-TBG-iGFP (GFP) or AAV8-TBG-iCre (Cre). **(B–C)** Western blot analysis of liver lysates from GFP and Cre mice. Each lane in (B) represents one mouse. **(C)** Quantification of phospho/total protein ratios. **(D–E)** Western blot analysis of primary hepatocytes from GFP and Cre mice stimulated with vehicle (basal), glucagon (200 nM, D), or insulin (20 nM, E) for the indicated times. Relative phosphorylation levels of pCREB^S133^ (D) and pAKT^S473^ (E) are shown as fold change over GFP at 0 min, normalized to total protein and indicated below each blot. Arrows in (D) highlight phospho-PKA substrates that were unchanged or increased in RACK1-deficient cells. **(F–G)** qPCR analysis of PKA target genes *G6PC* (F) and *PCK1* (G) in hepatocytes treated for 3 h with vehicle, insulin (20 nM), glucagon (100 nM), glucagon + insulin, cAMP (20 mM), or cAMP + insulin. **(H)** qPCR analysis of the indicated genes in liver tissues from GFP and Cre mice following a 6-hour fast. *, **, *** p <0.05, 0.01 and 0.001 vs GFP, respectively; ^#^ p <0.001 vs glucagon or cAMP; ^&^ p < 0.05 vs cAMP.

Consistent with this, RACK1 deficiency in primary hepatocytes reduced phosphorylation of most PKA substrates, although some were unaffected or even elevated. Additionally, glucagon-stimulated CREB phosphorylation was attenuated by RACK1 loss, whereas insulin-stimulated AKT phosphorylation remained unaltered (Figure 3D). Furthermore, RACK1 deficiency impaired glucagon-and cAMP-induced expression of gluconeogenic genes, such as G6PC and PEPCK1, but did not alter the insulin-mediated suppression of these genes (Figure 3F and G). This suppression of gluconeogenic gene expression was also confirmed in the livers of fasted mice lacking RACK1 (Figure 3H).

These findings demonstrate that RACK1 selectively regulates glucagon signaling by modulating PKA activity and downstream CREB phosphorylation, thereby influencing hepatic gluconeogenesis without affecting insulin signaling.

### RACK1 directly interacts with key components of the PKA signaling pathway

Given that RACK1 functions as a scaffolding protein [12, 13], we hypothesized that it may regulate PKA activity through direct interactions with PKA subunits, their regulators, and downstream effectors. To test this, we performed co-immunoprecipitation assays. To avoid potential interference from endogenous protein complexes that might mask RACK1’s binding sites to antibodies, we expressed Flag-tagged RACK1 in the livers of RACK1-deficient mice at levels below those of endogenous RACK1. Co-immunoprecipitation was then carried out using anti-Flag-conjugated Sepharose beads. As shown in Figure 4A, RACK1 selectively coimmunoprecipitated with both the RIIα and PKAcα subunits of PKA, GCGR, and CREB, but not with Gαs.

**Figure 4:**
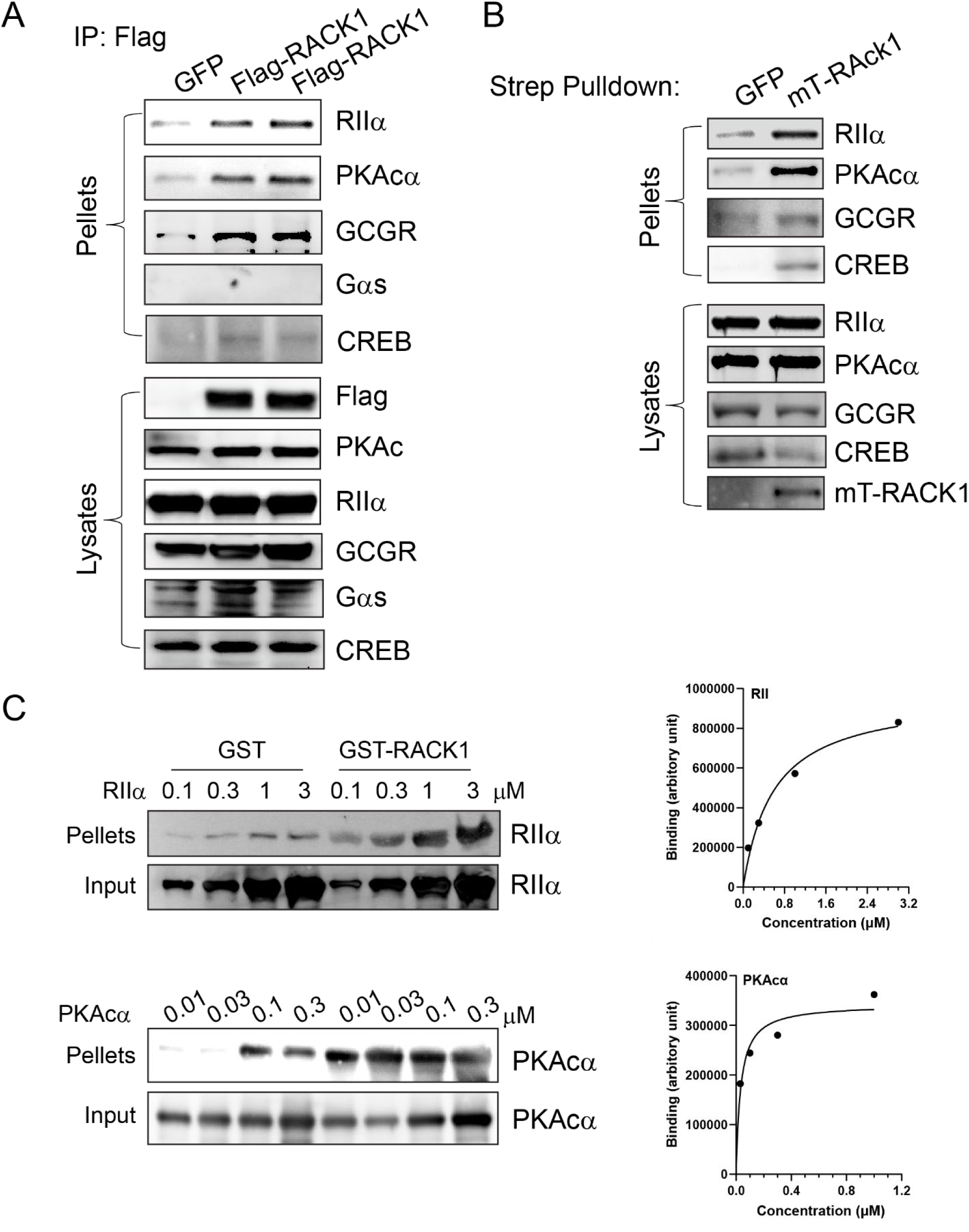
RACK1 interacts with PKA. **(A)** Co-immunoprecipitation analysis of liver lysates from Alb-Cre/RACK1^fl/fl^ mice injected with pAd-GFP or pAd-Flag-RACK1 demonstrates specific interactions between RACK1 and PKA subunits RIIα and PKAcα, as well as GCGR and CREB. **(B)** Streptavidin pull-down assays in liver lysates from Alb-Cre/RACK1^fl/fl^ mice mice expressing miniTurbo-tagged RACK1 (mT-RACK1) or GFP control (GFP) confirm proximity-based labeling of RACK1-associated proteins, including RIIα, PKAcα, GCGR, and CREB. **(C)** GST pull-down assays demonstrate direct, dose-dependent binding of RACK1 to PKA RIIα and PKAcα subunits. Left panel shows binding curve quantification.

To determine whether these interactions were direct, we expressed miniTurboID-tagged RACK1 in the livers of RACK1-deficient mice and administered biotin to allow proximity-dependent biotinylation. Streptavidin pulldown assays were then performed using tissue lysates prepared under denaturing conditions with RIPA buffer containing 1% SDS to eliminate indirect interactions. As shown in Figure 4B, RIIα, PKAcα, GCGR, and CREB were all biotinylated and enriched, confirming close proximity and supporting their direct association with RACK1.

To further validate the interactions between RACK1 and the PKA subunits RIIα and PKAcα, we performed in vitro GST pulldown assays using purified proteins. As shown in Figure 4C, GST-tagged RACK1 directly bound both RIIα and PKAcα. Quantitative analysis revealed that RACK1 had a significantly higher binding affinity for PKAcα (Kd = 0.032 ± 0.014 μM) than for RIIα (Kd = 0.588 ± 0.161 μM), indicating an approximately 18-fold stronger interaction with the catalytic subunit (Figures 4C).

To determine whether these interactions are regulated by glucagon stimulation, we performed co-immunoprecipitation assays in RACK1-deficient primary hepatocytes reconstituted with Flag-tagged RACK1. In unstimulated cells, RACK1 predominantly associated with RIIα, PKAcα, and GCGR, with minimal interaction observed with CREB (Figure 5A-B).

**Figure 5:**
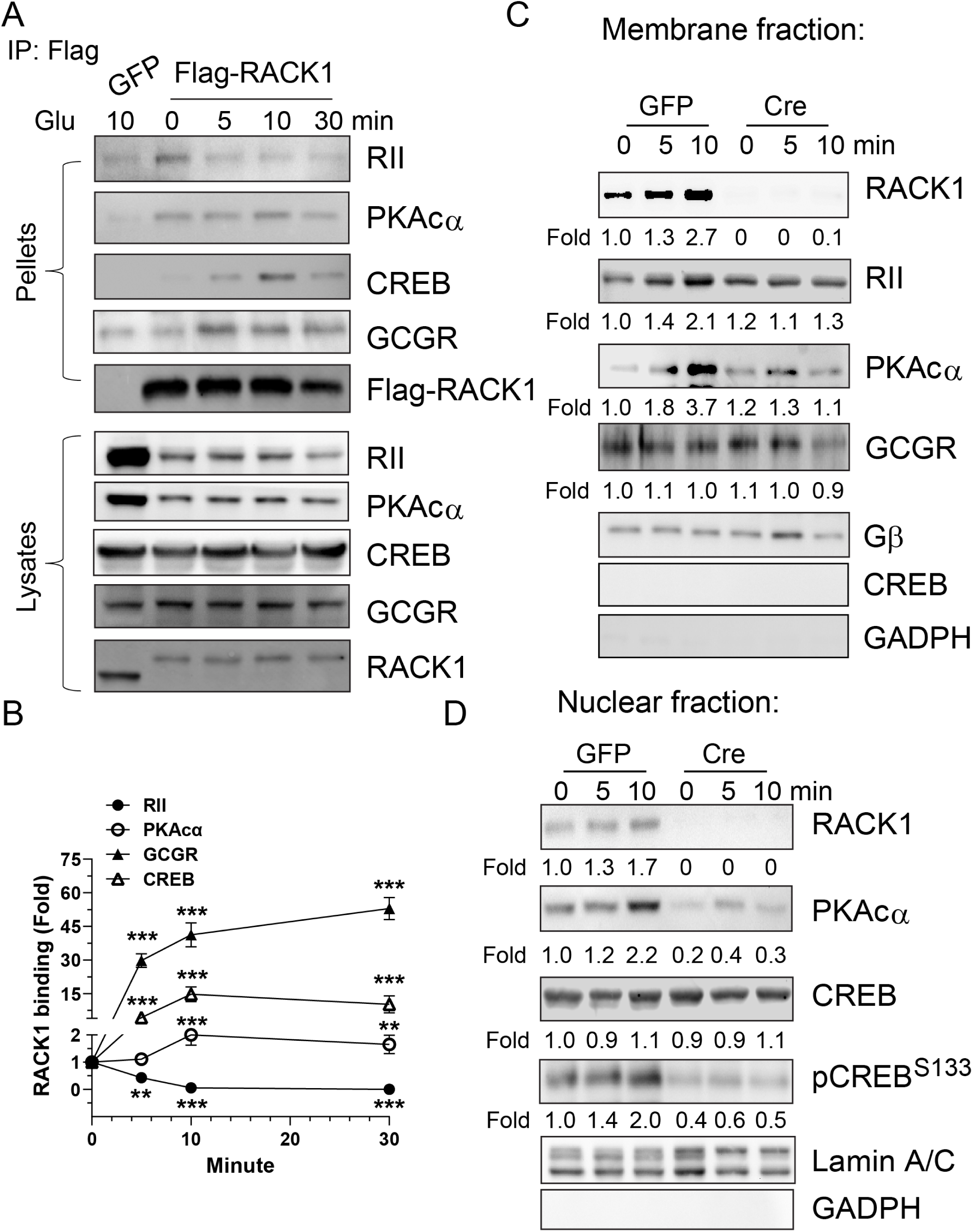
Glucagon regulates the dynamic interaction and compartmentation of RACK1 with components of the PKA signaling axis. **(A-B)** Co-immunoprecipitation of analysis of primary hepatocytes isolated from Alb-Cre/RACK1^fl/fl^ mice injected with pAd-GFP or pAd-Flag-RACK1 and treated with glucagon (200 nM) for the indicated times. Western blot analysis shows the time-dependent association of RACK1 with the PKA regulatory subunit RIIα, catalytic subunit PKAcα, GCGR, and CREB in IP pellets. Input blots (10% of lysates used for IP) confirm total protein expression levels. (**B)** Quantification of RACK1-associated proteins from (A), presented as fold enrichment relative to 0 min. **, *** p < 0.01 and 0.001 vs 0 min, respectively. N=3. **(C-D).** Western blot analysis of plasma membrane (C) and nuclear (D) fractions prepared from primary hepatocytes isolated from RACK1^fl/fl^ mice injected with AAV8-TBG-iGFP (GFP) or AAV8-TBG-iCre (Cre) and stimulated with glucagon (200 nM) for the indicated time. Glucagon-induced changes in protein localization were quantified as fold change relative to 0 min in GFP cells and are indicated below each blot.

Glucagon stimulation led to reduced interaction with RIIα but enhanced binding to PKAcα, GCGR, and CREB, suggesting that RACK1 preferentially associates with the PKA holoenzyme and GCGR under basal conditions, and shifts to interacting with GCGR, the dissociated PKA catalytic subunit and CREB upon GCGR and PKA activation.

Because GCGR and CREB are predominantly localized to membrane and nuclear compartments [10, 14], respectively, we investigated the spatial dynamics of these interactions. We performed cell fractionation of primary hepatocytes into membrane and nuclear compartments, with or without RACK1 deficiency (Figure 5C-D). Fraction purity was confirmed by the enrichment of membrane (G protein β subunits) and nuclear (lamin A/C) markers, and the absence of the cytosolic protein GAPDH. In unstimulated cells, small amounts of RACK1, and PKAcα were detected in the membrane fraction. Glucagon stimulation induced translocation of these proteins to the membrane, where GCGR is constitutively localized (Figure 5C). CREB, however, remained nuclear under both conditions (Figure 5D). In the nucleus, basal levels of RACK1, PKAcα, and phosphorylated CREB (pCREB^S133^) were present and further increased following glucagon stimulation. Importantly, RACK1 deficiency abolished glucagon-induced membrane translocation of PKAcα and significantly impaired nuclear accumulation of pCREB^S133^, indicating a role for RACK1 in facilitating spatial redistribution of PKA components and promoting CREB phosphorylation.

Confocal imaging further supported these observations. In unstimulated hepatocytes, RACK1 and PKAcα co-localized mainly in the cytosol and likely the membrane, with limited nuclear presence (Figure 6A-C). Upon glucagon stimulation, both proteins accumulated in the nucleus, without obvious enrichment in the membrane. Notably, RACK1 deficiency did not grossly alter PKAcα cellular distribution but completely prevented its nuclear translocation (Figure 6A-C), reinforcing RACK1’s critical role in nuclear delivery of PKAcα.

**Figure 6:**
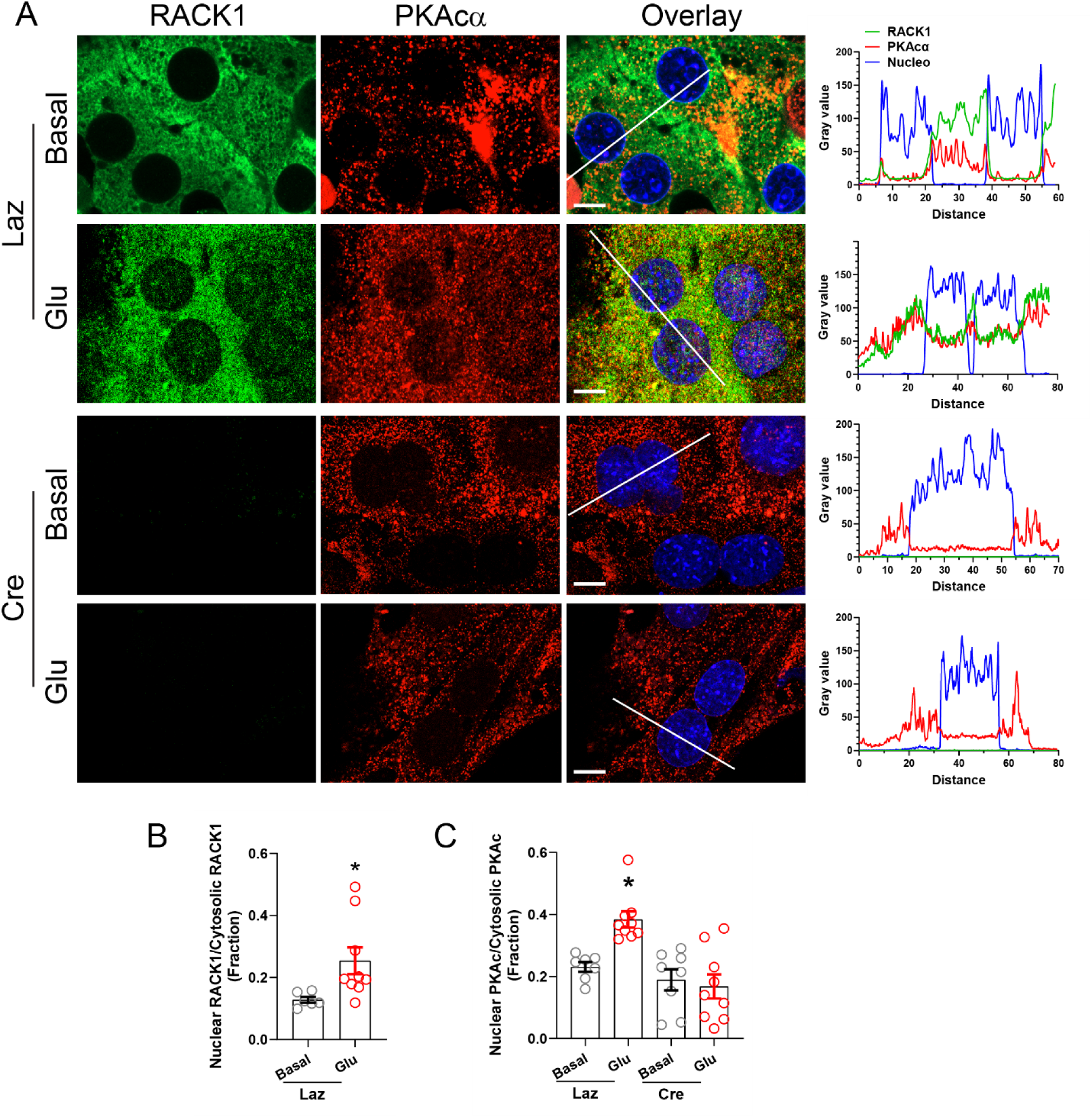
RACK1 is required for glucagon-induced nuclear translocation of PKAcα in primary hepatocytes. **(A).** Representative confocal immunofluorescence images of primary hepatocytes isolated from RACK1^fl/fl^ mice injected with adenovirus encoding either LacZ (Laz) or Cre (Cre), and cultured on collagen-coated coverslips. After overnight serum starvation, cells were stimulated with or without glucagon (100 nM) for 10 minutes, and stained with anti-RACK1 (green) and anti-PKAcα (red) antibodies. Nuclei were counterstained with DAPI (blue). Overlay panels show merged images. Line scans (white lines in overlay panels) with corresponding fluorescence intensity profiles for RACK1 (green), PKAcα (red), and nuclei (blue) are shown on the right. Scale bars, 20 μm. **(B-C).** Quantification of the nuclear-to-cytoplasmic fluorescence intensity ratio of RACK1 (B) and PKAcα (C), based on line scan analysis from confocal images. * p<0.05 vs basal.

In contrast, CREB was constitutively nuclear, regardless of glucagon stimulation or RACK1 expression (Figure 7A). However, glucagon promoted nuclear accumulation of RACK1 and enhanced co-localization with CREB (Figure 7A). The nuclear translocation of RACK1 appears essential for CREB phosphorylation, as RACK1 deficiency abolished CREB activation in both basal and stimulated conditions (Figure 7B-C).

**Figure 7:**
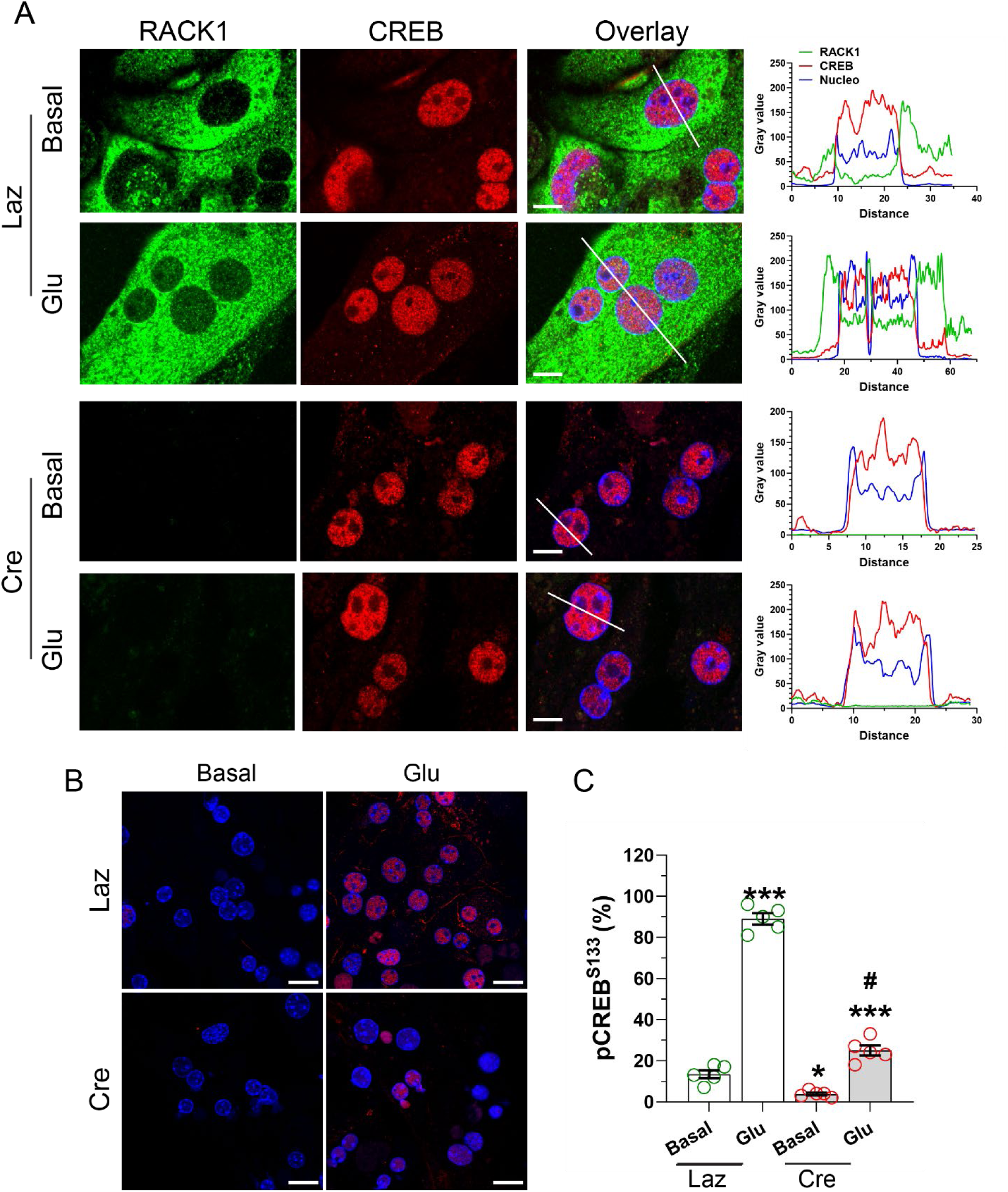
Nuclear translocation of RACK1 is required for glucagon-induced CREB phosphorylation in primary hepatocytes. **(A).** Representative confocal immunofluorescence images of primary hepatocytes prepared as in Figure 6 and stained with anti-RACK1 (green) and anti-CREB (red) antibodies. Nuclei were counterstained with DAPI (blue). Overlay panels show merged images. Line scans (white lines in overlay panels) with corresponding fluorescence intensity profiles for RACK1 (green), CREB (red), and nuclei (blue) are shown on the right. Scale bars, 20 μm. **(B-C).** Representative confocal immunofluorescence images of primary hepatocytes, prepared as in (A), and stained with anti-phospho-CREB^Ser133^ (red) antibody, and DAPI (blue). Scale bars, 10μm. (C) Quantification of nuclear pCREB ^Ser133^ signal is expressed as the percentage of pCREB-positive nuclei relative to total nuclei. *, *** p<0.05 and 0.001 vs basal, respectively. ^#^ p<0.001 vs glu in Laz.

Together, these findings demonstrate that RACK1 directly interacts with both regulatory and catalytic subunits of PKA, as well as upstream and downstream components including GCGR and CREB. These interactions are dynamically modulated by glucagon signaling and are required for proper compartmentalization and activation of the PKA pathway, culminating in CREB phosphorylation.

### RACK1 interacts with PKA and GCGR via the WD1-2 and WD3-4 domains to regulate PKA activation and gluconeogenesis

To investigate the molecular basis of RACK1’s interaction with PKA, GCGR and CREB, we generated a series of RACK1 deletion mutants corresponding to specific WD domains: WD1-2, WD3-4, and WD5-7 (Figure 8A). These mutants were GST-tagged and co-expressed with the RIIα and PKAcα subunits of PKA, GCGR or CREB in HEK293 cells. GST pulldown assays revealed that both RIIα and PKAcα, as well as GCGR preferentially interacted with the WD1-2 and WD3-4 domains. In contrast, CREB interacted broadly with all three domains, including WD1-2, WD3-4, and WD5-7 (Figure 8B).

**Figure 8.**
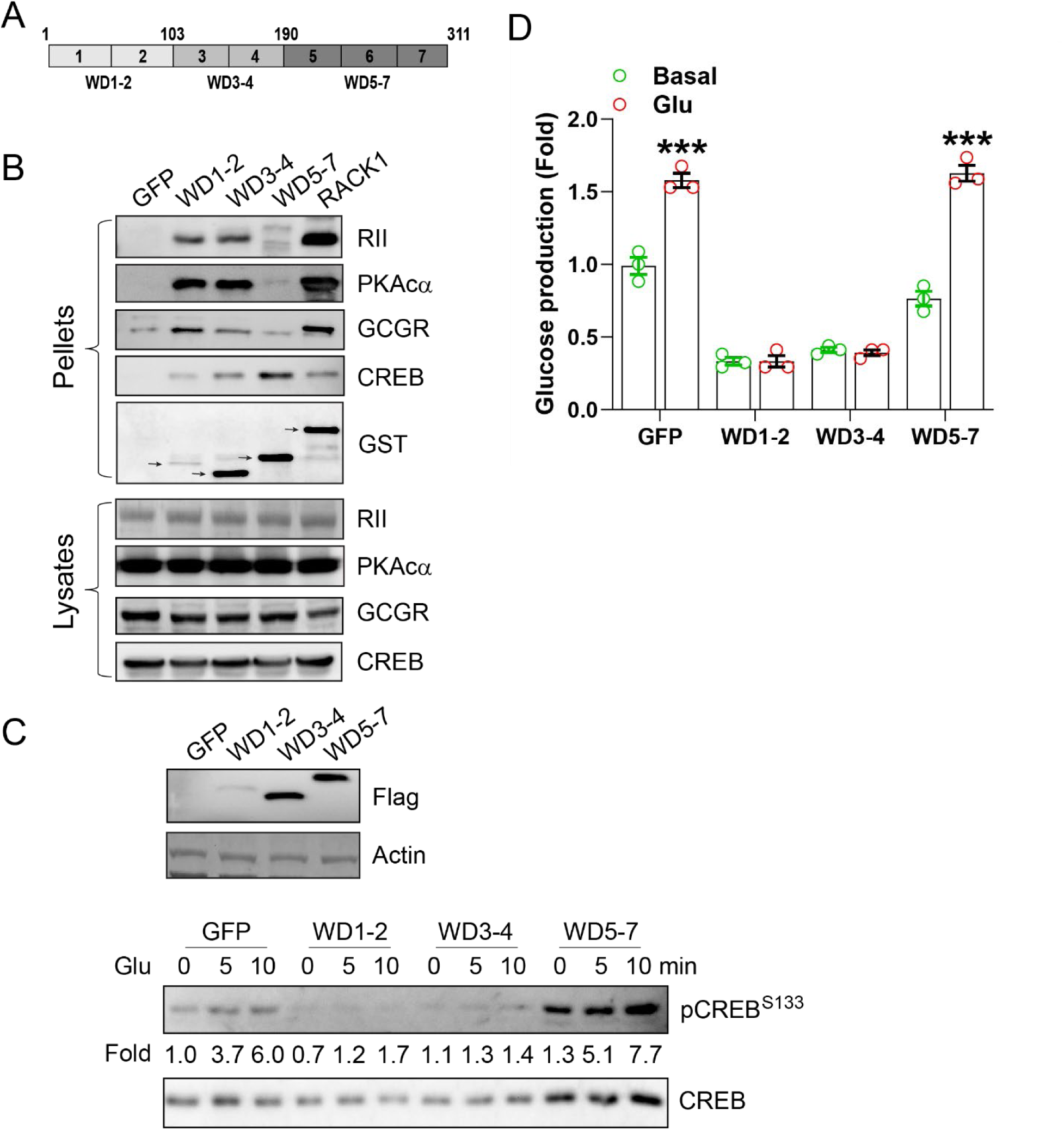
Molecular basis and functional role of RACK1 interactions with PKA, GCGR, and CREB. Schematic diagram of RACK1 deletion mutants containing various WD domains. The size of each construct is indicated. GST pull-down assays assessing interactions between RACK1 (full-length or mutants) and PKA RIIα, PKAcα, GCGR, and CREB. HEK293 cells were co-transfected with GFP control or GST-tagged RACK1 WD1-2, WD3-4, or WD5-7 constructs along with RIIα, PKAcα, GCGR, or CREB expression plasmids. Arrows indicate the positions of GST-tagged RACK1 and its domain mutants. **(C-D)** Functional impact of overexpressing Flag-tagged WD1-2, WD3-4, or WD5-7 domains on glucagon-induced CREB phosphorylation (C) and gluconeogenesis (D) in primary hepatocytes. Hepatocytes were infected with pAd-GFP (control) or Flag-tagged domain constructs. Western blot analysis shows relative levels of pCREB ^Ser133^ normalized to total CREB and expressed as fold change over GFP at 0 minutes. Expression of GFP and Flag-tagged mutants is shown in the top panel of C. ***p < 0.001 vs basal.

To assess the functional importance of these interactions, we overexpressed the RACK1 deletion mutants in primary hepatocytes and evaluated their effects on glucagon-stimulated PKA activation and glucose production. Overexpression of the WD1-2 and WD3-4 domains significantly suppressed glucagon-stimulated CREB phosphorylation (Figure 8C) and glucose production (Figure 8D). Notably, the WD1-2 domain exerted comparable inhibitory effects despite being expressed at lower levels than WD3-4, suggesting that WD1-2 may serve as the primary functional site mediating RACK1’s scaffolding activity. In contrast, overexpression of the WD5-7 domain—which does not interact with PKA or GCGR but binds CREB—had no significant effect on CREB phosphorylation or glucose output, even when expressed at levels similar to WD3-4 (Figure 8C-D).

These findings suggest that the WD1-2 and WD3-4 domains of RACK1 are critical for its interaction with PKA and GCGR and play an essential role in mediating glucagon-stimulated PKA activation and hepatic gluconeogenesis.

### Active PKAcα rescues defective glucose homeostasis induced by hepatic RACK1 deficiency

To determine whether the impaired glucose homeostasis observed in hepatic RACK1 deficiency results from suppressed PKA activation, we utilized transgenic mice expressing a constitutively active PKAcα mutant (PKAcα^W196R^) in a Cre-dependent manner [15]. Expression of the PKAcα^W196R^ mutant in the liver significantly increased the expression of gluconeogenic genes, including G6PC and PEPCK, compared to control mice (Figure 9A). Notably, while RACK1 deficiency alone suppressed gluconeogenic gene expression in control mice, this effect was completely abolished in mice expressing the PKAcα^W196R^ mutant, indicating that active PKAcα compensates for the loss of RACK1 (Figure 9A).

**Figure 9.**
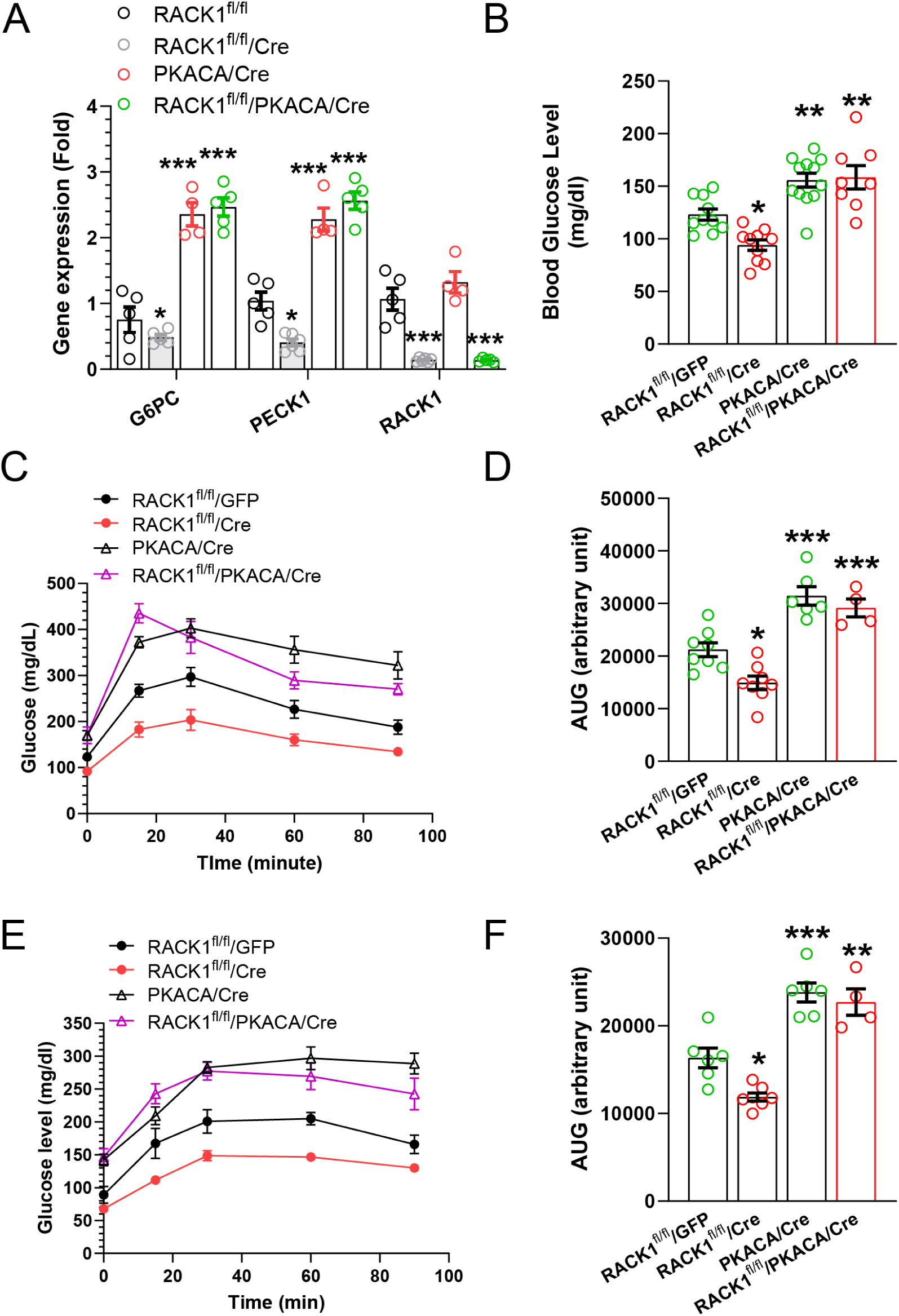
Rescue of glucose homeostasis defects induced by RACK1 deficiency via expression of constitutively active PKAc*α*^W196R^. **(A)** qPCR analysis of PKA target gene expression in livers of RACK1^fl/fl^ mice or RACK1-deficient mice (RACK1^fl/fl^/Cre) with or without expression of active ΠKAcα^W196R^ (PKACA/Cre). *, ***p < 0.05 and 0.001 vs RACK1^fl/fl^, respectively. **(B)** Basal blood glucose levels following a 24-hour fast. **(C–D)** Glucose tolerance test (C) and corresponding area under the curve (AUC) analysis (D). **(E–F)** Pyruvate tolerance test (E) and corresponding AUC analysis (F). *, **, ***p < 0.05, 0.01, and 0.001 vs. Cre (RACK1-deficient), respectively.

Consistent with these molecular findings, expression of the PKAcα^W196R^ mutant in the liver increased blood glucose levels in mice after 6 hours of fasting and reduced glucose and pyruvate tolerances during tolerance tests (Figure 9B-F). Importantly, combining hepatic RACK1 deficiency with the expression of the PKAcα^W196R^ mutant had no additional effects, and the glucose metabolism parameters were similar to those observed in mice expressing PKAcα^W196R^ alone (Figure 9B-F).

These results demonstrate that constitutively active PKAcα can overcome the defective gluconeogenesis and glucose homeostasis caused by hepatic RACK1 deficiency, further underscoring the pivotal role of RACK1 in regulating PKA-mediated glucose metabolism.

## Discussion

Previous research has highlighted the significance of the glucagon-PKA signaling axis in glucose metabolism [11, 16], but the mechanisms governing the spatial and functional coordination of this pathway have remained incompletely understood. Our study reveals a previously unrecognized and essential role for RACK1 in maintaining hepatic glucose homeostasis by regulating glucagon-PKA-CREB signaling. Acute, liver-specific RACK1 deficiency leads to profound metabolic consequences, including fasting hypoglycemia, impaired glucose tolerance, and diminished hepatic glucose production. These findings align with prior reports showing that chronic hepatic RACK1 deficiency disrupts glucose homeostasis [17], further supporting a direct and critical role for RACK1 in the regulation of hepatic glucose metabolism.

Mechanistically, RACK1 loss selectively impairs glucagon-stimulated, but not insulin-mediated, signaling pathways, resulting in markedly reduced phosphorylation of CREB and suppressed expression of key gluconeogenic genes such as *G6PC* and *PCK1*. Notably, this defect occurs without changes in hepatic cAMP levels, indicating that RACK1 does not influence glucagon receptor signaling upstream of cAMP production but rather acts at the level of PKA activation and downstream effector targeting.

The critical importance of PKA signaling in this phenotype is underscored by rescue experiments using a constitutively active PKAcα mutant (PKAcα^W196R^), which restores gluconeogenic gene expression, fasting blood glucose levels, glucose and pyruvate tolerance in RACK1-deficient mice. These findings directly link RACK1 to the regulation of hepatic gluconeogenesis through modulation of PKA activity, and exclude broader roles of RACK1 in transcriptional control or metabolism under acute conditions.

Our findings also uncover a unique mechanistic function of RACK1 as a spatial organizer of PKA signaling in hepatic gluconeogenesis. While PKA activity is classically regulated by A-kinase anchoring proteins (AKAPs) [9], which tether PKA to specific subcellular compartments via its regulatory subunits most liver-expressed AKAPs have limited impact on glucagon-stimulated gluconeogenesis. One exception is radixin, a plasma membrane–anchored AKAP that modulates glucagon-PKA signaling through its interaction with the membrane protein prom1 [10]. In contrast to traditional AKAPs, RACK1 directly binds both the regulatory (RIIα) and catalytic (PKAcα) subunits of PKA, as well as the upstream receptor GCGR and downstream transcription factor CREB. Importantly, RACK1 exhibits stronger binding affinity to PKAcα than RIIα, distinguishing it from typical AKAPs that interact exclusively with regulatory subunits. This suggests that RACK1 may play dual roles—anchoring the inactive PKA holoenzyme to GCGR through the regulatory subunits and facilitating the cellular specific localization of activated PKAcα for downstream signal propagation.

Supporting this model, overexpression of RACK1 WD1–2 or WD3–4 domain, which mediate interactions with GCGR, PKA, and CREB and likely acts in a dominant negative manner to disrupt these interactions, attenuates glucagon-induced CREB phosphorylation and gluconeogenesis. In contrast, overexpression of the WD5–7 domain, which interacts only with CREB, has no effect. These findings underscore the importance of RACK1 simultaneously engaging multiple components of the glucagon–PKA–CREB pathway to coordinate effective signal propagation. Cell fractionation and co-immunoprecipitation studies further demonstrate that under basal conditions, RACK1 forms a membrane-associated complex with GCGR, RIIα, and PKAcα. Upon glucagon stimulation, RACK1 is required for recruiting PKA to the membrane for effective assembly with GCGR. Interestingly, stimulation reduces RACK1–RIIα interaction but enhances its association with PKAcα and CREB, consistent with PKA activation and subunit dissociation. Although confocal imaging reveals cytosolic co-localization of RACK1 and PKAcα, no marked translocation to the membrane is observed, suggesting only a small, signaling-competent pool is involved. These findings underscore the need for more sensitive tools to visualize dynamic RACK1–PKA interactions in live cells and further dissect their spatial regulation of glucagon signaling.

Our confocal imaging and subcellular fractionation analyses demonstrate that RACK1 is essential for the nuclear translocation of PKAcα and subsequent phosphorylation of CREB in hepatocytes. In the absence of RACK1, glucagon stimulation fails to induce nuclear accumulation of PKAcα or activate CREB, despite normal intracellular cAMP levels— underscoring RACK1’s pivotal role in bridging PKA activation to nuclear transcriptional responses and gluconeogenesis.

The mechanisms driving glucagon-induced translocation of RACK1 to the plasma membrane and nucleus remain to be elucidated. Notably, earlier studies in hippocampal neurons have shown that activation of the cAMP/PKA pathway can promote nuclear translocation of RACK1 [18]. Whether similar regulatory mechanisms are engaged downstream of glucagon signaling in hepatocytes remains to be determined.

This dual-compartment scaffolding model, in which RACK1 organizes GCGR–PKA complexes at the plasma membrane and PKA–CREB complexes in the nucleus, provides a mechanistic framework for how spatial precision is conferred upon the glucagon-PKA-CREB signaling axis. By facilitating the dynamic assembly and translocation of PKA signaling components, RACK1 may ensure efficient and selective transcriptional activation of gluconeogenic genes. To our knowledge, this is the first report to identify RACK1 as a spatial coordinator of the glucagon-PKA-CREB signaling pathway. While prior studies have implicated RACK1 in PKA regulation through interactions with proteins such as PDE4D5, which modulate cAMP degradation [19, 20], our findings uncover a distinct mechanism whereby RACK1 directly scaffolds key signaling components to enable compartmentalized signal transduction.

In addition to its role in regulating hepatic glucose homeostasis, glucagon signaling also plays a central role in hepatic lipid metabolism by suppressing de novo lipogenesis [11, 21]. Previous studies using chronic hepatic RACK1 deficiency models driven by Alb-Cre have shown that long-term loss of RACK1 leads to lipid accumulation, hepatic steatosis, and ultimately tumorigenesis [17, 22]. However, it remains unclear whether these phenotypes result directly from RACK1 loss or from progressive metabolic compensation. Future studies are needed to assess the acute effects of RACK1 deficiency on lipid metabolism to distinguish primary from secondary consequences.

Notably, glucagon may suppresse hepatic lipogenesis via mechanisms distinct from those regulating gluconeogenesis, primarily through the inhibition of the transcription factor SREBP1c (Sterol Regulatory Element-Binding Protein 1c) and activation of AMPK, both mediated by the direct or indirect action of PKA [21, 23, 24]. Our findings suggest that RACK1 deficiency differentially affects PKA substrates. While glucagon-stimulated phosphorylation of CREB is markedly suppressed in the absence of RACK1, other PKA substrates display variable responses (Figure 3D). These observations suggest that RACK1 may contribute to substrate specificity within the PKA signaling cascade, enabling differential regulation of glucose and lipid metabolism. Elucidating how RACK1 confers this specificity may provide novel insights into the spatial and functional compartmentalization of PKA signaling in the liver.

In conclusion, our findings reveal a novel and mechanistically distinct role for RACK1 in coordinating PKA signaling to control hepatic glucose metabolism. RACK1 scaffolds both regulatory and catalytic subunits of PKA with GCGR at the membrane to initiate signaling and facilitates the nuclear translocation of PKAcα for downstream CREB activation. This dual-compartment scaffolding function enables spatial and substrate-specific control of PKA activity, which is critical for regulating gluconeogenesis during fasting. These insights not only clarify the functional organization of the glucagon-PKA-CREB axis but also suggest that RACK1 could be a promising therapeutic target for the treatment of metabolic disorders such as type 2 diabetes, where excessive hepatic glucose production contributes to hyperglycemia [3]. Future studies exploring the regulation of RACK1 under metabolic stress and its broader impact on hepatic lipid metabolism may further expand its relevance in metabolic disease pathogenesis and therapy.

## MATERIALS AND METHODS

### Reagents

Glucagon and insulin were obtained from Millipore Sigma (St. Louis, MO, USA). H89 and compound 3i (3i) were purchased from Shelleck Chemicals (Houston, TX, USA). Antibodies for CREB (no. 9197), phospho-CREB^S133^ (no. 9198), AKT (no. 4685), phospho-AKTS473^S473^ (no. 4060), and PKAcα (no. 4782) were procured from Cell Signaling Technology (Danvers, MA, USA); GADPH (sc-47724) from Santa Cruz Biotechnology (Dallas, TX, USA); mouse anti-RACK1 (no. 61077), RIIα (no. 612242), and PKAc (no. 610980) from BD Biosciences (Franklin Lakes, NJ, USA). Rabbit anti glucagon receptor (GCGR; no. 26784-1-AP) was purchased from Proteintech (Rosemont, IL, USA). Horseradish peroxidases-conjugated secondary antibodies against mouse and rabbit IgG were from Jackson ImmunoResearch (West Grove, PA, USA).

Alexa 448-and 568-conjugated anti mouse or rabbilt IgG were from Thermo Fisher Scientific (Waltham, MA, USA). Mouse anti-Flag-conjugated Sepharose beads were purchased from BioLegend (San Diego, CA, USA). Streptavidin-magnetic beads were obtained from Pierce (Thermo Fisher Scientific, Waltham, MA, USA).

### Animal studies

Animal studies were performed in accordance with an IACUC-approved protocol (#3011932) at the University of Iowa. C57BL/6J mice with Loxp-flanked exon 2 and 3 in the RACK1 gene (RACK1^fl/fl^) were utilized, as reported previously [25]. Deletion of RACK1 in the liver in fetal mice was achieved by crossing with transgenic mice expressing Cre recombinase driving by the albumin promoter and enhancer (Alb-Cre; #035593; Jackson Laboratory, Bar Harbor, ME, USA) to produce AlbCre/RACK1^fl/fl^ mice. Acute deletion of RACK1 gene in the liver was achieved through the injection of AAV viruses (1×10^11^ genome copies/mouse) encoding Cre recombinase under the thyroxine-binding globulin (TBG) promoter (AAV-TBG-iCre; Vector Biolabs, Malvern, PA, USA) into the retroorbital vein of RACK1^fl/fl^ transgenic mice. Mice were studied 10–14 days post-injection.

Transgenic mice (#032825) carrying a Cre-dependent expression of a constitutively active PKAcα mutant W196R (PKAcα^W196R^) were purchased from the Mutant Mouse Regional Resource Centers and backcrossed with C57BL/6J mice for at least nine generations. Genotyping was performed using a combination of PCR and restriction enzyme Mlu I digestion, as described previously [15]. Liver-specific expression of PKAcα^W196R^ was induced by injecting AAV-TBG-iCre, as described above.

Blood glucose levels were measured from tail tips using a glucometer (Accu-Chek, Roche Diabetes Care Inc., USA). For glucose and pyruvate tolerance tests, 12-16-week old male mice were fasted for 18 hours with access to water *ad libitum* and intraperitoneally injected with D-glucose (1.5 g/kg body weight) or a pyruvate/lactate mixture at a 1:10 weight ratio (1.0 g/kg body weight). For the insulin tolerance test, male mice were fasted for 6 hours and intraperitoneally injected with insulin (1.0 U/kg body weight).

### Isolation of mouse primary hepatocytes

Primary hepatocytes were isolated from 12−16 week-old male and female mice using liberase (Sigma-Aldrich, St. Louis, MO, USA), following a previously established protocol [26]. The isolated hepatocytes were suspended in William’s E medium supplemented with 10% fetal bovine serum (FBS), 10 nM dexamethasone, and 20 nM insulin. Cells were plated on collagen-coated dishes for subsequent experiments.

### Glucose production assays

Glucose production in mouse primary hepatocytes was measured as described previously [27]. Briefly, hepatocytes cultured on collagen-coated plates were serum-starved overnight in glucose-free DMEM. The cells were then incubated with glucose production buffer containing glucose-free, phenol red-free DMEM supplemented with 20 mM sodium lactate, 2 mM sodium pyruvate, 2 mM L-glutamine, and 15 mM HEPES. Treatments included glucagon (200 nM) in the presence or absence of insulin (20 nM) or other inhibitors. After 18 hours, glucose levels in the culture media were quantified using a colorimetric glucose detection kit (Thermo Fisher Scientific, USA), according to the manufacturer’s instructions.

### cAMP measurement

The cAMP levels in liver tissues from fasted mice were quantified using the mouse/rat cAMP Parameter Assay Kit (Bio-Techne, Minneapolis, MN, USA) [28]. Concentrations were normalized to protein content, which was determined using a BCA protein assay.

### Western blotting analysis

Protein lysates were prepared from cultured cells and tumor tissues and analyzed via Western blotting as previously described [25, 28].

### Quantitative real-time PCR (qPCR)

Total RNA was extracted from liver tissues and primary hepatocytes using the Zymo Direct-zol RNA Mini Preparation Kit (Zymo Research, Irvine, CA, USA). RNA was used to analyze target gene expression by qPCR with the SensiFAST SYBR No-ROX Kit (Bioline USA Inc., Taunton, MA, USA), following previously established protocols [25]. The primer sequences used for qPCR were: G6PC (forward: 5’-tcttgtggttgggattctgg, reverse: 5’-cggatgtggctgaaagtttc), PEPCK1 (forward: 5’ - ccatcccaactcgagattctg, reverse: 5’ ctgagggcttcatagacaagg), RACK1 (forward: 5’ - aatactctgggtgtctgcaag, reverse: 5’ - ttagccagattccacaccttg).

### Plasmids and DNA constructs

RACK1 and its deletion mutants (WD1-2, WD3-4, and WD5-7) were previously constructed in the pENTR/SD entry vector [29]. These constructs were cloned into the pDEST27 destination vector to express GST-tagged proteins in mammalian cells using the Gateway cloning system (Thermo Fisher Scientific, USA).

Flag-tagged RACK1 and its deletion mutants were generated by adding a Flag tag through PCR amplification. To create miniTurbo-tagged RACK1, the RACK1 sequence was first cloned into the pcDNA3-C1(1-29)-miniTurbo-V5 vector (#107174, Addgene, Watertown, MA, USA). The miniTurbo-tagged RACK1 sequence was then amplified by PCR and cloned into the pENTR/SD vector. Subsequently, these constructs were transferred into the pAd-CMV-DEST destination vector for adenoviral production.

### Adenoviral production and transduction

Adenoviruses encoding Flag-RACK1, its deletion mutants, and miniTurbo-RACK1 were generated by transfecting PacI-digested pAd-CMV-DEST vectors into HEK293A cells. Viruses were amplified in HEK293A cells and purified by ultracentrifugation using a CsCl_2_ density gradient.

For in vivo expression, adenoviruses (1 × 10^11^ pfu/mouse) were administered via retroorbital injection into 8–10-week-old AlbCre/RACK1^fl/fl^ mice to drive liver-specific expression of GFP, Flag-RACK1, or miniTurbo-RACK1. For in vitro expression in primary hepatocytes, cells were incubated overnight with adenoviruses at a multiplicity of infection of ∼50 pfu/cell.

### Co-immunoprecipitation assay

Liver lysates were prepared in modified RIPA buffer containing 50 mM Tris-HCl (pH 7.4), 150 mM NaCl, 1 mM EDTA, and 1% Triton X-100. The lysates were incubated with mouse anti-Flag antibody-conjugated Sepharose beads for 6 hours at 4°C. Co-immunoprecipitated proteins were then analyzed as described previously [25].

### Purification of recombinant proteins

GST-fusion RACK1 was expressed in *E. coli* BL21 cells as described previously [30]. Cell pellets were lysed by sonication in a buffer containing 100 mM Tris-HCl (pH 7.4), 200 mM NaCl, 1 mM EDTA, 1% NP-40, and 1 mM DTT. The lysates were centrifuged at 100,000 × g for 1 hour at 4°C, and 40% glycerol was added to the supernatant. The resulting protein preparations were stored at-20°C.

His-tagged RIIα and PKAcα proteins were expressed in *E. coli* BL21 cells following induction with 0.5 mM IPTG overnight at 18°C [31, 32]. The proteins were purified to >90% purity using nickel affinity chromatography and stored in a HEPES buffer containing 10 mM HEPES (pH 7.2), 50 mM NaCl, 10 mM MgCl_2_, and 40% glycerol at-20°C.

### GST pulldown assays

For GST pulldown assays using purified proteins, 5 μg of GST or GST-RACK1 from the bacterial lysates was isolated with 50 μl of a 50% slurry of Glutathione-Sepharose beads (Gold Biotechnology, St. Louis, MO, USA). These beads were incubated with purified RIIα and PKAcα proteins in 100 μl of binding buffer (100 mM Tris-HCl, pH 7.4, 200 mM NaCl, 1 mM DTT, 0.1% Triton X-100, and 10 mM imidazole) for 30 minutes at room temperature [33]. The beads were washed five times with ice-cold RIPA buffer. The bound proteins were resolved by SDS-PAGE and analyzed via Western blotting.

To analyze GST-RACK1 and its mutant binding to PKA, GCGR and CREB in HEK293A cells, cells were transiently transfected with plasmids encoding GFP, GST-RACK1, or its mutants along with pcDNA3-RIIα-V5 and PKAcα-V5, GCGR or CREB using PolyJet transfection reagent (SignaGen Laboratories, Frederick, MD, USA) in 100 mm plates. Two days post-transfection, cell lysates were prepared using RIPA buffer and incubated with 60 μl of a 50% slurry of Glutathione-Sepharose beads for 1 hour at room temperature. The pulled-down proteins were resolved by SDS-PAGE and analyzed via Western blotting.

### Streptavidin pulldown assay

7 days after AlbCre/RACK1^fl/fl^ mice were injected with adenoviruses encoding GFP or miniTurbo-RACK1, the mice were intraperitoneally injected with biotin (30 μg in 100 μl of PBS) and either maintained on a normal diet or fasted for 24 hours [34]. Liver lysates were then prepared from excised liver tissue using RIPA buffer. 3 mg of the lysates were incubated with 50 μl of streptavidin-magnetic beads for 4 hours at 4°C. After washing five times with RIPA buffer, the bound proteins were analyzed by Western blotting.

### Cell fractionation

Plasma membrane and nuclear fractions were isolated from primary hepatocytes using a modified subcellular fractionation protocol [25, 35]. Briefly, cells were homogenized on ice using a hand-held pestle homogenizer in 50 mM Tris-HCl (pH 8.0) buffer containing 0.5 mM DTT, 0.1% NP-40, and protease and phosphatase inhibitor cocktails. The homogenates were first centrifuged at 750 × g for 10 minutes at 4°C to pellet plasma membrane-enriched fractions, and centrifuged at 10,000 × g for 10 seconds to collect the nuclear pellet. Fraction purity was confirmed by immunoblotting: the absence of cytosolic GAPDH and the enrichment of G protein β subunits and lamin A/C were used to validate membrane and nuclear fractions, respectively.

### Immunofluorescence staining and Confocal imaging Analysis

Primary hepatocytes isolated from RACK1^fl/fl^ mice injected with adenovirus encoding either LacZ (control) or Cre recombinase and cultured on collagen-coated coverslips. After overnight serum starvation, cells were stimulated with or without glucagon (100 nM) for 10 minutes. Cells were then fixed with 4% paraformaldehyde and permeabilized using 0.25% Triton X-100. Following permeabilization, cells were incubated overnight at 4°C with primary antibodies: rabbit anti-PKAcα (1:100), rabbit anti-CREB (1:100), or rabbit anti-phospho-CREB (Ser133) (1:100), in combination with mouse anti-RACK1 (1:100), as we reported [25]. The next day, cells were incubated with Alexa Fluor 488- or 568-conjugated secondary antibodies (1:800; anti-rabbit or anti-mouse IgG) for 1 hour at room temperature. Nuclei were counterstained with DAPI.

Z-stack images were captured using a Carl Zeiss LSM800 Meta inverted confocal microscope equipped with a 63× oil immersion objective lens (NA 1.3). Image analysis and quantification were performed using Fiji (ImageJ).

### Statistics

Data were expressed as mean ± SEM. Statistical comparisons between groups were analyzed by two tail Student’s *t* test or ANOVA with *P*<0.05 considered statistically significant.

## Abbreviations

AKAP: A-kinase anchoring protein
cAMP: cyclic AMP
CREB: cAMP response element-binding protein
GCGR: glucagon receptor
G6PC: glucose-6-phosphatase catalytic subunit
GFP: green fluorescent protein
GST: glutathione S-transferase
PECK: Phosphoenolpyruvate carboxykinase 1
PGC1: peroxisome Proliferator-activated receptor gamma coactivator 1-alpha
PKA: protein kinase A
RACK1: receptor for activate C kinase 1
WD: ryptophan-aspartate repeat Domain

## ACKNOWLEDGMENTS

We would like to thank Maddi Lensing, and William Draus for their assistance in maintaining and breeding the transgenic mice. This work was supported in part by funding from National Institutes of Health grant R01CA282699 (SC) and pilot grant from Fraternal Order of Eagle Diabetes Research Center, University of Iowa.

## AUTHOR CONTRTIBUTIONS

L.C. and S.C. performed the experiments and data analysis; L.C. and S.C. designed the experiments with inputs from L.Y.; S.C. wrote the manuscript and L.Y. provided critical review of the manuscript. S.C. obtained funding for the study.

